# A Deep Learning Framework for Spatiotemporal Modeling of Visual Task fMRI

**DOI:** 10.64898/2026.05.05.722117

**Authors:** Mingyang Li, Yiwei Chen, Chengling Ning, Xixi Dang, Dan Wu

## Abstract

Characterizing the dynamic coordination of distributed brain regions during cognitive tasks remains challenging, as traditional fMRI analysis focuses on localized activations without revealing the underlying information flow that drives them. Here, we propose STREAM (Spatiotemporal Representation for Effective connectivity Analysis Model), a deep-learning framework that learns neural transition functions in task-fMRI to characterize effective connectivity and whole-brain information flow. Applied to visual category processing in 1074 participants, STREAM accurately reconstructs activation maps while further revealing that traditional activation regions are primarily driven by incoming signals. Moreover, the Default Mode Network acts as a high-level regulatory hub with extensive outgoing influence, challenging its passive characterization. Additionally, category-specific communication emerges from dynamic reconfiguration of signaling patterns among key hubs rather than static pathways. These findings establish a novel computational paradigm that uncovers directional signaling mechanisms driving local dynamics in task-fMRI, revealing how the brain flexibly reconfigures functional architecture for complex cognition.

## Introduction

The advances of cognitive neuroscience has been profoundly shaped by research using blood-oxygen-level-dependent (BOLD) based task-fMRI, which provides unprecedented insights into the neural basis of diverse cognitive domains^1^, including vision^2^, language^3^, and working memory^4^. The predominant paradigm in task-fMRI studies relies on the General Linear Model (GLM) to identify brain regions specialized for particular cognitive processes. In the field of visual neuroscience, for instance, this localization-based approach has successfully identified a mosaic of category-selective regions within the ventral visual stream^2,5^, such as the fusiform face area (FFA) for face perception^6^, the parahippocampal place area (PPA) for scene recognition^7^, and distinct clusters for bodies and tools^8,9^. Furthermore, large-scale mapping has revealed that these regions are embedded within a hierarchical organization, progressing from elementary feature extraction in early visual areas toward higher-order semantic integration in prefrontal and parietal networks^10–13^.

While activation analysis provides a robust “where” perspective by pinpointing cognitive hubs, it primarily captures the local metabolic endpoints of task-induced activity, remaining fundamentally silent on the dynamic signaling flow that drives these responses^14^. The directed information flux, or effective connectivity (EC), represents the underlying mechanism by which the brain’s functional architecture is dynamically reconfigured, serving as the critical link between broad task demands and localized neural dynamics^15^. To capture the directional orchestration of the visual processes, high-temporal-resolution techniques like magnetoencephalography (MEG) and stereo-electroencephalography (sEEG) have identified crucial recurrent interactions along the ventral stream^16–21^. However, these methods often suffer from limited spatial depth and source ambiguity, failing to pinpoint the precise whole-brain topography of these interactions. Complementary evidence from dynamic causal modeling (DCM) and vector autoregression (VAR) have been developed to formally infer directed signaling from fMRI data^22,23^. These models have shown that visual category processing, while driven by bottom-up inputs, is flexibly modulated by top-down feedback^24,25^. However, most of these fMRI-based models rely on rigid, linear assumptions that fail to capture the brain’s inherent non-linear neural dynamics^26^, and require pre-selecting a limited set of regions of interest (ROIs), typically informed by GLM activation maps^27,28^. Given that GLM activation is thought to reflect signal convergence and post-synaptic integration^29,30^, such approaches may bias toward information-receiving hubs and systematically obscure the primary sources of outgoing flow. Specifically, we reason that task-evoked activations are not isolated events but localized manifestations of a brain-wide signaling reconfiguration. In this view, distinct task demands first trigger the specific up- or down-regulation of directed information flow to dynamically orchestrate distributed brain regions into task-specific functional networks. Within this process, the directional signaling exchange at each node define the local dynamic profiles, where in-flow and out-flow may play distinct roles in shaping local neural activations. Therefore, fully deciphering the neural mechanisms of cognition necessitates the reconstruction of this spatiotemporal signaling process, as these directed interactions constitute the fundamental causal backbone that both ignites localized responses and orchestrates distributed regions into a coherent functional system.

The emergence of deep learning neural networks offers a transformative paradigm to bridge these methodological gaps. By modeling fMRI time series as continuous trajectories of a dynamical system, these architectures learn the latent neural transition functions embedded within high-dimensional data^22,28,31^, from which directed interactions can be causally inferred^32–34^. A recent study introduced the Neural Perturbational Inference (NPI) framework, demonstrating that virtual perturbations on a deep-learning-based brain surrogate can infer whole-brain effective connectivity from resting-state fMRI, outperforming traditional methods like DCM^28^. Grounded in the fundamental principle of learning neural transition functions, we assume perturbational framework could be extended to task fMRI to characterize directed information flow during cognitive processing.

Here, we introduce Spatiotemporal Representation for Effective connectivity Analysis Model (STREAM), a deep learning framework designed to infer directed information flow from task-evoked neural dynamics. We first desgined the framework architecture by identifying the optimal network structure, and the applied STREAM to characterize visual category processing within a large-scale cohort from the Human Connectome Project (HCP). As illustrated in Figure 1, we first demonstrate that STREAM can accurately capture and predict task-based neural dynamics, providing a robust foundation for causal inference. Based on this framework, we decomposed directed information flow into In-flow and Out-flow components. While the In-flow component is robustly coupled with localized activation maps, the Out-flow component is largely decoupled from these maps but reconciles with the default network. Furthermore, STREAM identifies a common recurrent hierarchical backbone shared across all categories, as we specialized processing emerges through a dynamic reconfiguration of directed signaling, where key neural hubs flexibly reroute their communication patterns to meet unique task demands. By resolving the localized activations into their underlying signaling drivers, this work reveals the dynamic reconfiguration of brain-wide information flow that sustains diverse visual processing tasks, establishing a scalable computational paradigm that can be generalizable to a broad range of cognitive domains.

**Figure 1.**
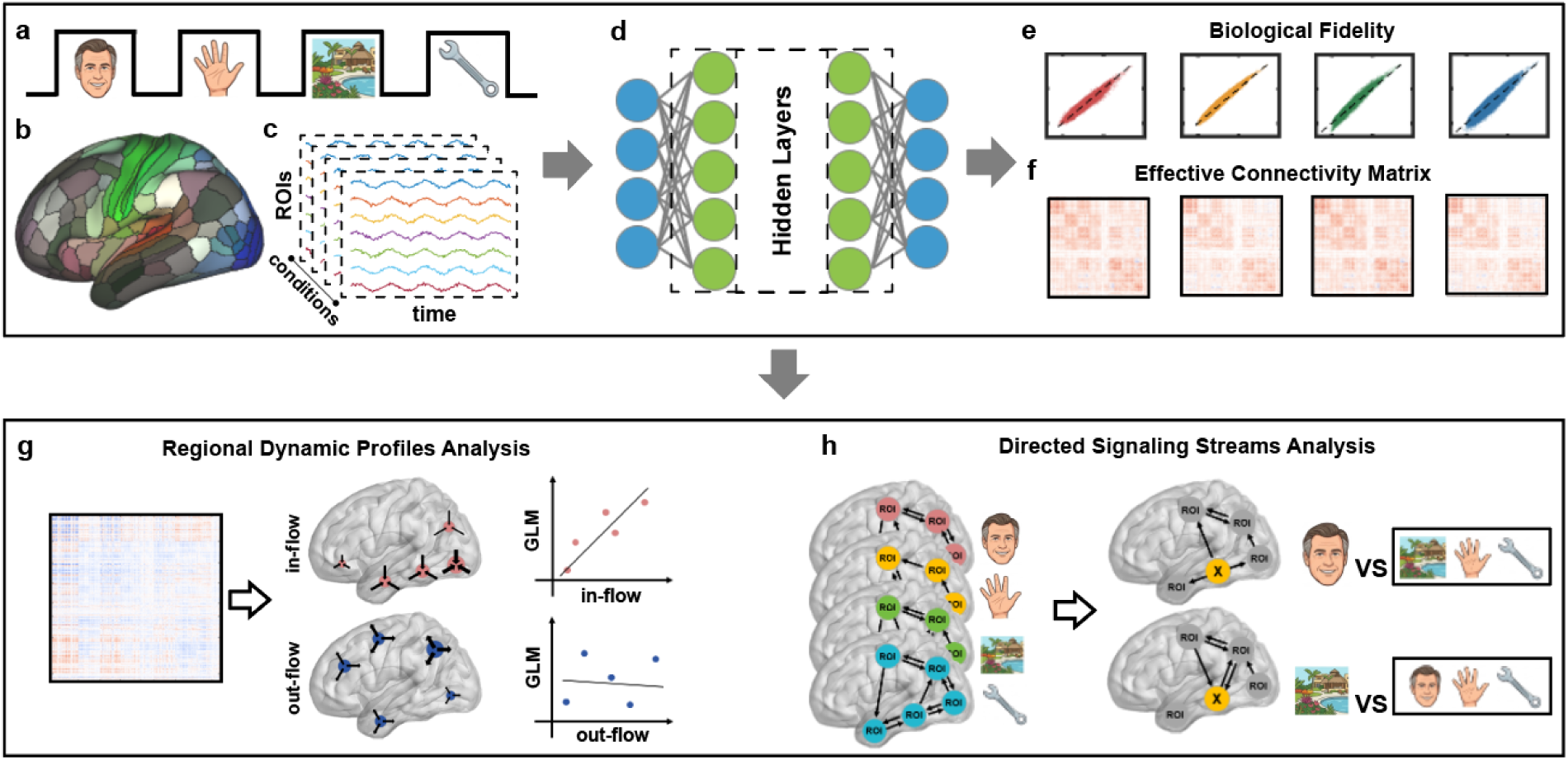
Overview of the research pipeline. (a) A block-design visual task featuring four stimulus categories (Faces, Body, Places, and Tools). (b–c) BOLD time series were extracted from 379 ROIs encompassing the whole brain. (d) A deep neural network architecture designed to capture the non-linear transition functions and underlying temporal trajectories of brain states from empirical fMRI data. (e) The model’s biological validity was assessed by comparing simulated signals with empirical fMRI data. (f) Whole-brain effective connectivity (EC) was obtained from the trained network via Neural Perturbation Inference, measuring directional causal influences between ROI pairs by simulating virtual perturbations. (g) Regional dynamic signatures were defined by decomposing EC matrices into in-flow and out-flow components, which were then spatially correlated with GLM-based activation maps. (h) Signaling streams were identified by contrasting one condition against the others (e.g., Face vs. Others), analogous to GLM contrasts, to isolate category-selective information routing.

## Results

### Design of the STREAM framework

To find a neural network architecture best for vision task-MRI, we evaluated five types of networks, Multilayer Perceptron (MLP), Recurrent Neural Network (RNN), Vector Autoregression (VAR), Long Short-Term Memory (LSTM), and Residual Network (Residual). The networks were trained using a segment of existing fMRI signals to predict the signal at the next time point, as an initial setting based on established benchmarks in previous literature^28^. We used a history length of three time points as the input to forecast the subsequent volume. Models were trained on each individual independently and evaluated across 100 participants and five task conditions (see Methods). For each model, the time series from each task condition was temporally split, with the first 80% of time points used for training and the remaining 20% held out for testing, preserving the sequential structure of the BOLD signals. All models demonstrated robust convergence, with training and test loss curves stabilizing within 100 epochs. To assess biological fidelity, we used the trained models to iteratively generate synthetic BOLD signals and compared their functional connectivity (FC) topology to the empirical task FC via Pearson correlation. Under this evaluation, the MLP architecture demonstrated high fidelity by faithfully reconstructing the empirical FC (*r* = 0.935, rank 3/5; Figure S1 and Table S1), while simultaneously achieving a low prediction error (MSE = 0.719 ± 0.213, rank 2/5). While the VAR and Residual models achieved slightly higher FC correlations (*r* = 0.983 and 0.981, respectively), their predictive accuracy was comparatively lower (MSE = 0.868 ± 0.248 and 0.737 ± 0.213). Notably, despite its high correlation coefficient, the VAR model exhibited systematic bias (Figure S1). Overall, the Residual and MLP models showed equivalent performance. Considering its architectural simplicity and superior efficiency, with an average runtime per individual-condition model (2.62 ± 0.76 s) nearly 3-fold faster than the Residual architecture (7.49 ±1.61 s), we selected the MLP as the core of the STREAM framework.

We further investigated the influence of the length of fMRI signals used for prediction (1-5 time points) on the MLP’s performance (Figure S2 and Table S2). While the node-level predictive accuracy was highest at signal length of 1 (MSE = 0.692 ± 0.182) and gradually declined as the length extended, the reconstruction of the group-level FC topology improved significantly with longer steps, rising from 0.886 at length=1 to 0.949 at length=5. To balance temporal prediction accuracy and functional connectivity fidelity, a signal length of 3 was identified as the optimal for next-time-point prediction. Building upon this selection, we further refined the MLP architecture by increasing the network depth to three hidden layers and expanding the hidden layer width to ensure more stable convergence and enhanced generalization. This optimized STREAM framework was then applied to the subsequent systematic analysis.

### High-Fidelity and Robust Learning of Task-Based Neural Dynamics

We first evaluated the prediction performance of STREAM and its robustness across the entire cohorts (n=1074) for the five experimental conditions, including Face, Body, Place, Tool, and Baseline. The training and test loss curves for all conditions demonstrated a rapid convergence (Figure 2a), suggesting that the MLP successfully learned the underlying neural dynamics without overfitting. Regarding the reconstruction of functional architecture, the model captured feasible FC profiles across all five conditions at the individual level. The mean individual FC correlation values ranged from 0.539 to 0.701 across conditions, suggesting a moderate performance in reconstructing individual-level FC topology (Figure 2b), similar to previous studies using rsfMRI^28^. At the group level, the simulated FC matrices exhibited extremely high similarity with the empirical group-average, with all correlation coefficients exceeding 0.93 (ranging from 0.932 to 0.982; Figure 2c-d, Table S3). These results demonstrate that by aggregating across subjects to mitigate individual noise, STREAM reliably reconstructs the accurate functional architecture at task level.

**Figure 2.**
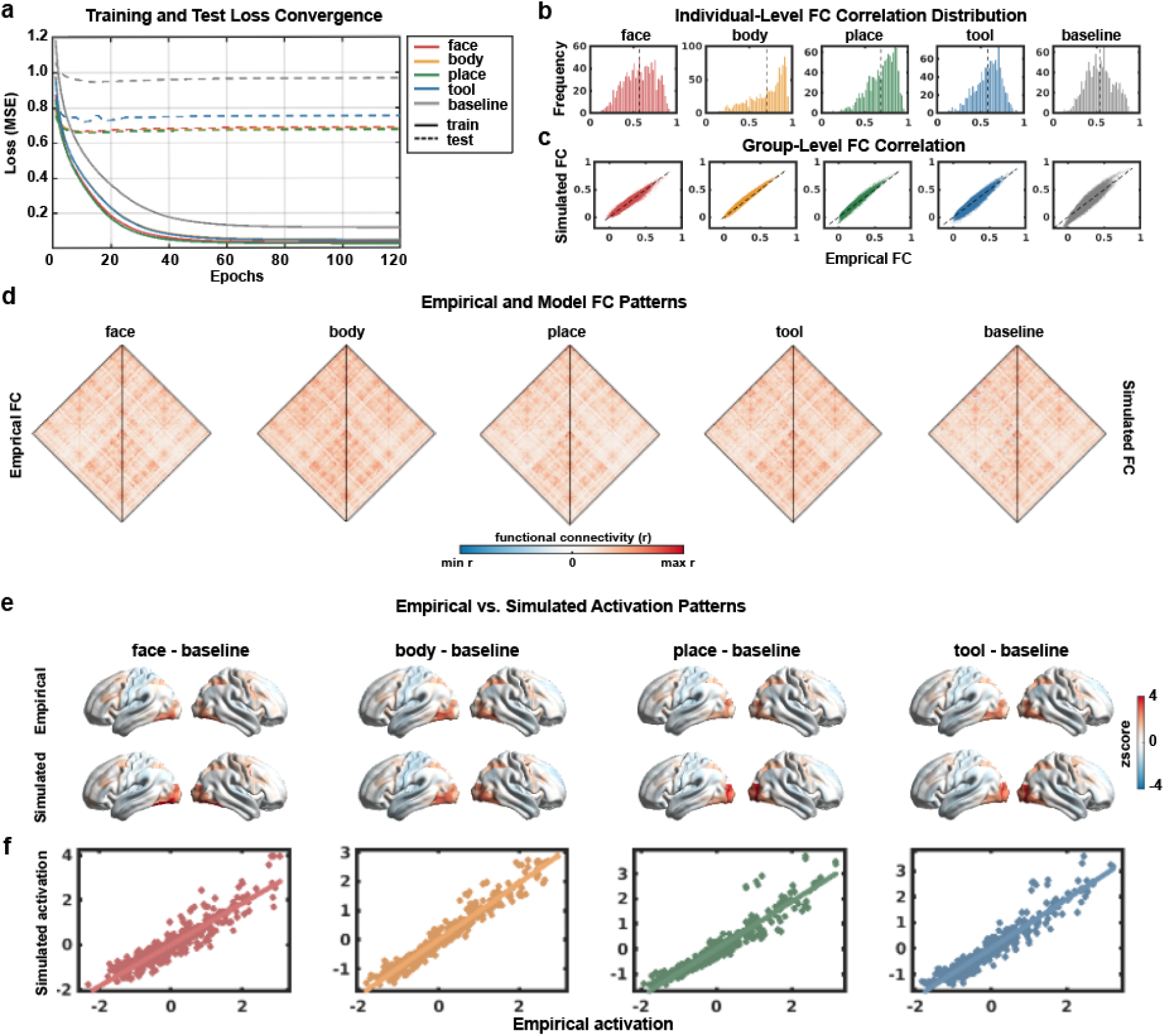
High-fidelity reconstruction of task-based neural dynamics and activation patterns. (a) Training (solid) and test (dashed) loss curves across the five experimental conditions, demonstrating stable convergence within 120 epochs. (b) Histograms showing the distribution of Pearson correlation coefficients between empirical and simulated functional connectivity (FC) for each participant across five conditions. (c) Scatter plots illustrating the robust correspondence between empirical and simulated group-averaged FC across all conditions (r > 0.93). (d) Diamond plots display empirical FC (left) and simulated FC (right) for each condition. Note that colormaps are individually scaled to their respective ranges (min–max r) to facilitate the comparison of topological structures. (e) Surface renderings show group-level activation maps (Condition vs. Baseline) for empirical GLM (top) and simulated profiles (bottom). (f) Scatter plots showing the high spatial correlation between empirical Cohen’s d maps and simulated activation profiles across 379 regions (all *r* > 0.92).

Beyond connectivity, we examined whether simulated signals from STREAM could reflect localized activation patterns. We generated activation profiles of the simulated signals (Condition minus Baseline) and compared them with empirical group-level Cohen’s d maps. Both empirical GLM and simulated activity were predominantly localized within the visual cortex (Figure 2e). Quantitatively, the spatial correlation between simulated profiles and GLM maps across 379 regions exceeded 0.92 for all four tasks (Figure 2f and Table S3). Finally, to ensure the generalizability of these findings, we performed a split-half reliability analysis, which resulted in over 0.98 consistency across all conditions (Figure S3 and Table S3).

### General Organization of Effective Connectivity during Visual Processing

Based on the validated STREAM framework, we characterized the global EC patterns underlying visual cognitive tasks. EC was inferred using the NPI approach^28^, which quantifies directed influence by measuring the causal response of the trained deep-learning surrogate to systematic virtual perturbations (see Methods). While individual task conditions were analyzed separately, they exhibited a consistent spatial organization (see Figure 3 for Face and Place; Figures S4-S6 for Body, Tool, and Baseline). In all conditions, positive ECs predominantly defined the connectivity landscape, accounting for over 99% of significant edges (e.g., 87698 positive vs. 688 negative ECs in the Face condition; Table S4). We then dissected the overall nodal strength into In-flow and Out-flow components and mapped them onto the cortical surface. Figure 3c-d revealed a common hierarchical gradient among all task conditions: positive In-flow and Out-flow were predominantly centered within the visual cortex and exhibited a graded organization along the ventral stream. In contrast, regions with strong negative In-flow and Out-flow were primarily distributed across frontal, parietal, and temporal cortices, largely outside the visual system.

**Figure 3.**
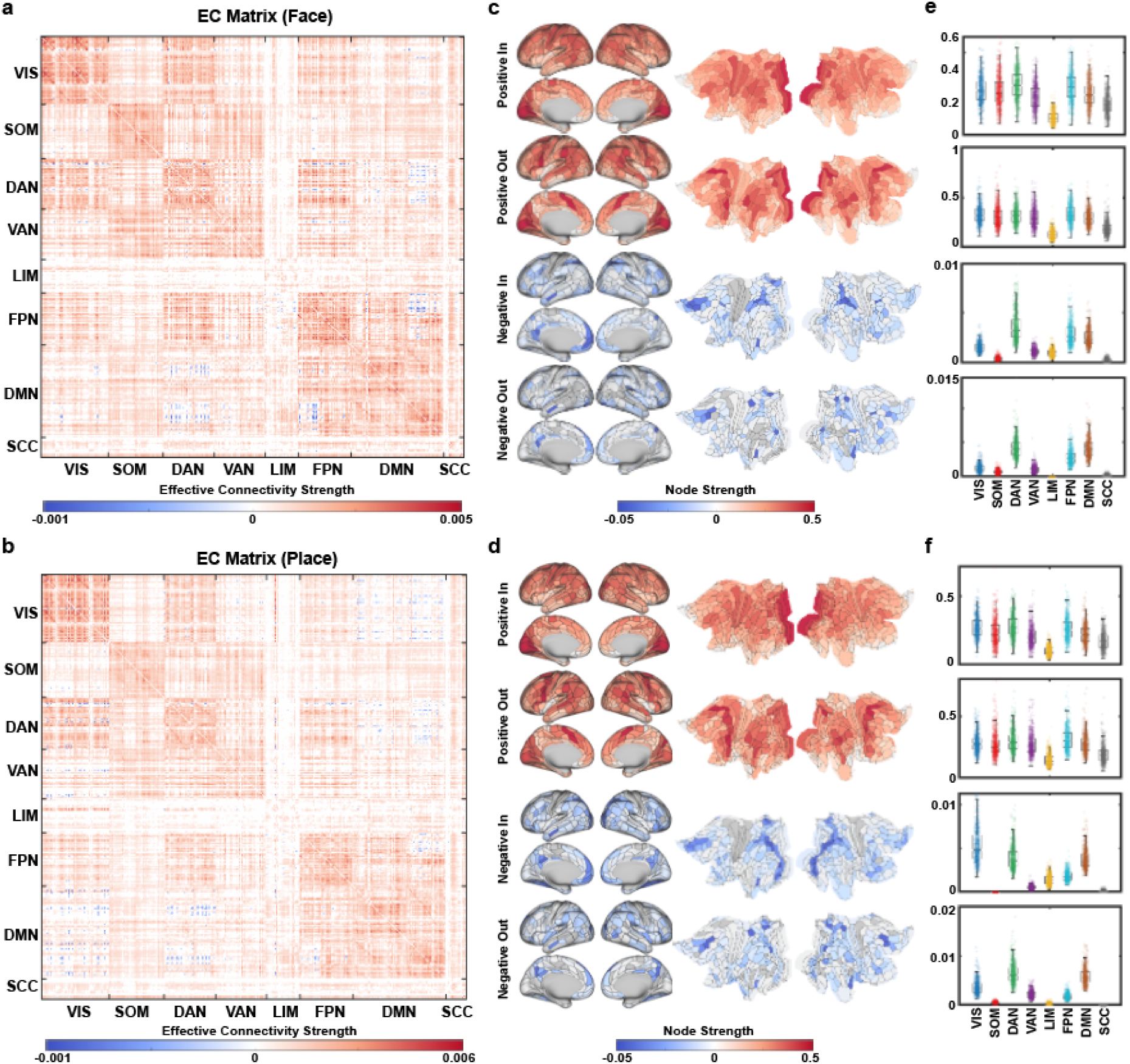
Global organization of effective connectivity during visual tasks. (a–b) Matrices illustrate the significant effective connectivity (EC) for (a) Face and (b) Place conditions (Bonferroni corrected *p* < 0.05). Regions are ordered according to seven canonical functional networks and one subcortical module. (c–d) Surface and flattened map renderings for (c) Face and (d) Place conditions, displaying the spatial distribution of four flow metrics (positive In-flow, positive Out-flow, negative In-flow, and negative Out-flow). Each row presents the standard cortical view (left) and the corresponding flattened surface (right), highlighting the hierarchical gradient centered in the visual cortex. (e–f) Violin plots for (e) Face and (f) Place conditions quantify the nodal flow across the eight functional modules. Each panel corresponds to the four flow metrics shown in (c–d), with individual data points representing participants and the violin structure illustrating the cross-subject distribution within each canonical network.

To further quantify these common features, we partitioned the EC results into seven canonical functional networks^35^ and one subcortical module (Figure 3e-f). Across all five experimental conditions, the positive ECs exhibited a widespread distribution that recruited all subnetworks, highlighting the broad coordination between sensory and high-level cognitive systems during task engagement. In contrast, negative ECs—representing inhibitory interactions—showed a more localized and sparser organization. These inhibitory flows were primarily concentrated within the Visual (VIS), Dorsal Attention (DAN), and Default Mode (DMN) networks. This pattern may reflect a fundamental regulatory mechanism in visual tasks, where sensory regions receive task-relevant input while being dynamically modulated by suppressive signals from higher-order association cortices to balance external processing with internal cognitive demands.

The stability of this shared EC architecture was confirmed by split-half reproducibility test that showed high correlation (*r* > 0.97 for all conditions, Figure S7) and high spatial consistency of the significant edges (DICE coefficient > 0.92; Table S4).

### Task-Driven Regulation of Effective Connectivity and Its Coupling with Local Activation

Beyond the general EC backbone, we hypothesize that distinct cognitive demands trigger brain-wide signaling reconfigurations, where task-evoked regulations of directed information flow converge to define local dynamic signatures. To test this, we compared these nodal signatures with traditional GLM-based activation maps using differential EC matrices (ΔEC_baseline,_, i.e., regulation of the baseline EC, Table S5), derived by subtracting baseline from each task condition (Figure 4a and 4e for Face and Place; see Figure S8 for Body and Tool). Nodal regulation intensity was quantified by summing up- and down-regulated in-flow and out-flow for each region. These intensities revealed that in-flow regulations were predominantly concentrated in the ventral occipitotemporal cortex (VOTC) (Figure 4b, f). Spatial correlation analysis showed that across all categories, positive GLM activations were robustly coupled with up-regulated in-flow (*r*: 0.71–0.84) and significantly correlated with down-regulated in-flow (*r*: −0.57 to −0.30) (Figure 4d, h and Figure S8). This finding suggests that the BOLD signal reflects the total synaptic processing load within a target region, regardless of whether the net effect is enhanced or suppressed input. When restricting the spatial correlation analysis to these statistically significant edges, the coupling between GLM activation and EC regulation became even more pronounced (Figure S9).

**Figure 4.**
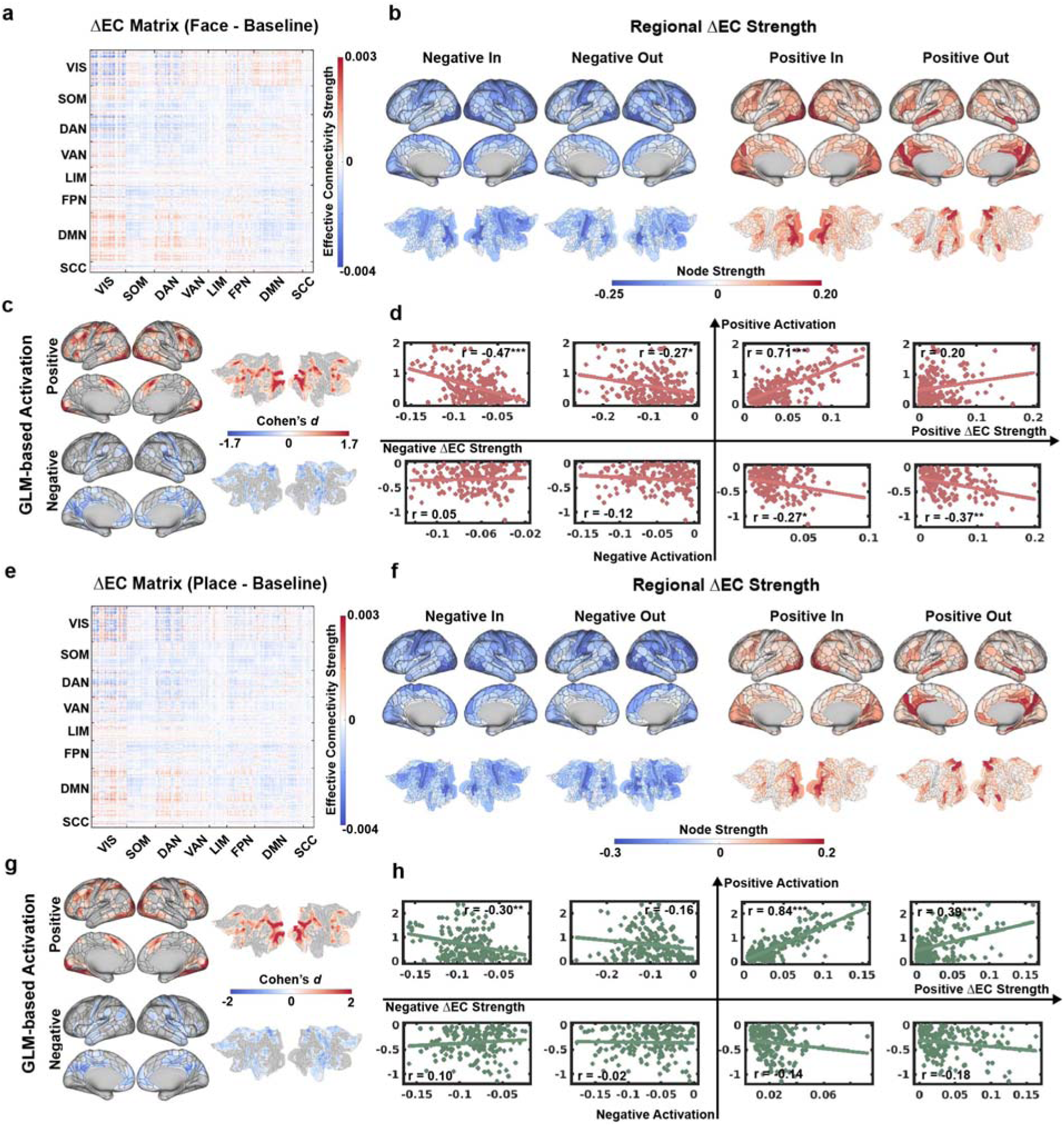
Task-Driven Regulation of Effective Connectivity and Its Coupling with Local Activation. (a, e) Differential EC Matrices (ΔEC). Matrices for Face (a) and Place (e) conditions show unthresholded task-induced regulation (Condition vs. Baseline; red: up-regulation; blue: down-regulation), with regions organized by functional networks. (b, f) Cortical Topography of EC Regulation. Surface and flattened maps visualize four regulation dimensions: Positive/Negative In-flow and Out-flow. (c, g) Local Task Activation. Group-level Cohen’s d maps showing positive and negative BOLD activations (GLM) for both conditions. (d, h) Spatial Coupling. Scatter plots quantify the spatial correspondence between GLM activation (Positive/Negative) and the four EC regulation metrics. The plots are arranged in four quadrants corresponding to the GLM polarity and EC regulation direction. P-values were corrected for multiple comparisons using Bonferroni correction. *p < 0.05, ** p < 10^−3^, *** p < 10^−6^.

At the systems level (Figure S10), up-regulated in-flow peaked in the VIS, DAN, and Frontoparietal Network (FPN) networks (all t > 79, p < 0.001), reflecting a massive influx into sensory and attentional hubs. Conversely, up-regulated out-flow was concentrated in the DMN and FPN (t > 88, p < 0.001), identifying these higher-order networks as primary sources of regulatory signals. Intriguingly, these out-flow hubs showed limited spatial correlation with GLM maps, revealing a fundamental dissociation: traditional activation mapping predominantly captures the convergence of inputs (in-flow) rather than the broadcast of influence (out-flow). Down-regulations were more widespread, with the strongest suppression in VIS, DAN, and Somatomotor (SOM) networks (all t > 78, p < 0.001), indicating task engagement involves broad inhibition of both incoming and outgoing signaling.

To test the generalizability of this coupling, we computed ΔEC_others_ matrices for category-selective contrasts (e.g., Face vs. Others, Table S6). We consistently found that positive GLM activations robustly correlated with up-regulated in-flow, while showing only weak correlations with out-flow (Figures S11–S13). Interestingly, for category-specific contrasts, negative GLM activations also showed strong coupling with down-regulated in-flow—a pattern less evident in task-versus-baseline comparisons. This discrepancy likely arises because the baseline condition involves meaningful task-related cognitive activity rather than an unconstrained resting state.

In summary, these results demonstrate that task-evoked activations are embedded within a global signaling reconfiguration. While the convergence of incoming signals is concentrated in visual processing areas and robustly coupled with GLM activations, the broadcast of outgoing influence is distributed across the DMN and FPN, potentially mediating top-down attentional control and task-related modulation.

### Recurrent Hierarchical Information Flow across the Ventral Stream

Having established that traditional GLM activations primarily reflect the convergence of incoming signaling (In-flow), we next sought to characterize the dynamic routes through which information reaches these functional hubs. While the brain maintains a stable structural scaffold, the functional accessibility of specific category-selective regions would be dynamically reconfigured to prioritize task-relevant information.

To provide a topographic perspective, we calculated the effective path cost from L-V1 to all other ventral visual areas (Figure 5). At the global level, the information flow across all five conditions followed a highly consistent recurrent hierarchical cascade, originating in L-V1 and progressing through a shared sequence of early visual areas including V2 and V3 (Figure 5 and Figure S14). However, as the signal reached the higher areas, the processing paths began to diverge according to the task demand. For instance, during the Place condition, the VMV and PHA exhibited a shorter cost from V1 compared to the Face condition, indicating higher information accessibility. In contrast, during the Face task, the FFC was positioned closer to the V1 area than it was during the Place task. This flow pattern suggests that while the visual system employs a universal feature-extraction mechanism in its early stages, the recurrent hierarchical architecture is dynamically reconfigured at higher-order stages to prioritize task-relevant information. Similar patterns were observed when using the R-V1 as the seed region.

**Figure 5.**
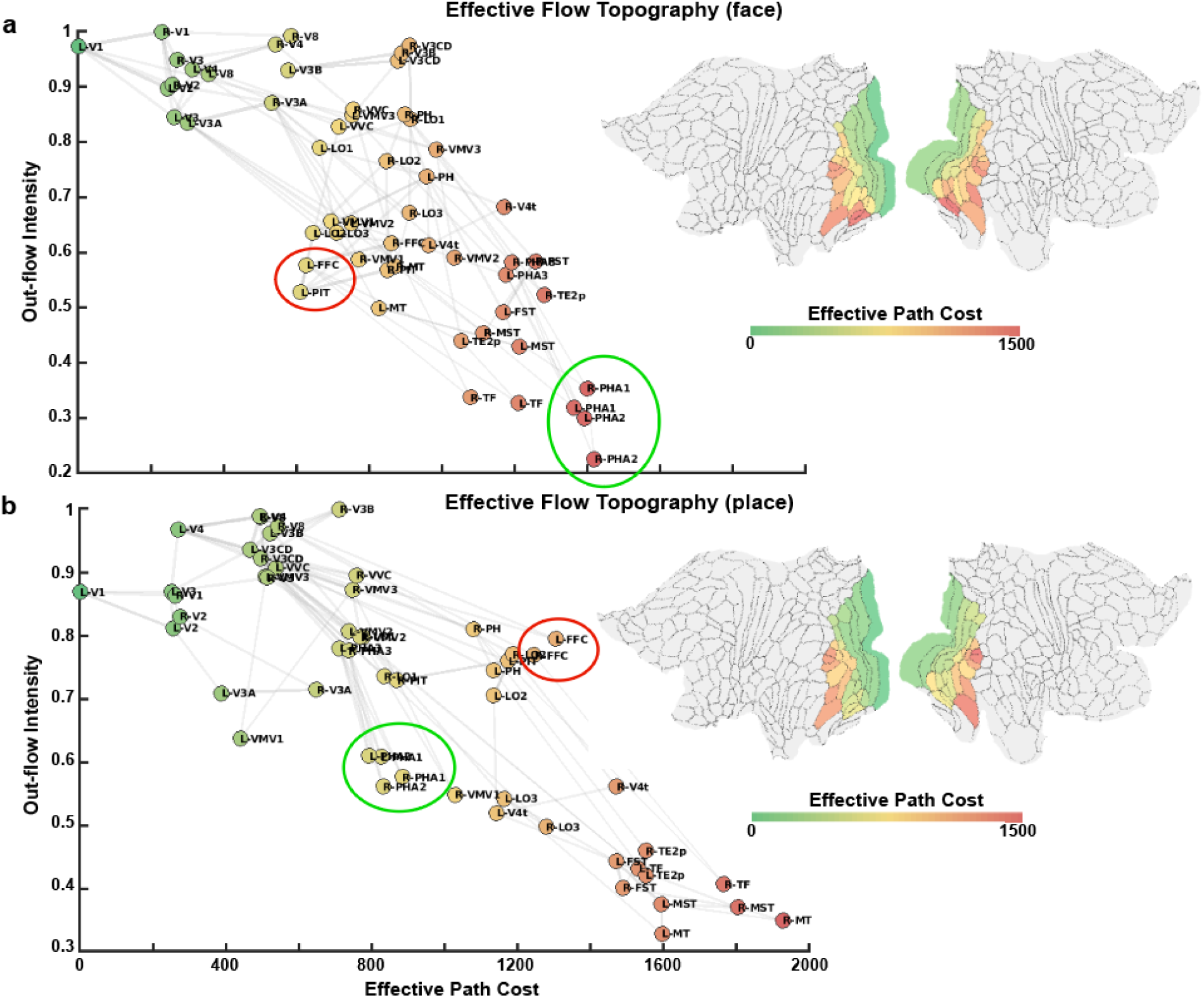
Recurrent Hierarchical Information Flow across the Ventral Stream. The plots visualize the hierarchical organization for (a) Face and (b) Place conditions. The x-axis represents the distance (effective path cost from L-V1), and the y-axis denotes the normalized out-flow intensity. Red circles highlight the face-relevant cluster (e.g., FFC), which shifts to a lower cost (closer to V1) during the face task. Green circles highlight the place-relevant cluster (e.g., PHA), showing increased accessibility during the place task. Insets show the flattened cortical representation of the ventral visual stream, with the colormap indicating the effective path cost from L-V1.

### Task-Specific Reconfiguration of Signaling Patterns in Category-Selective Hubs

The above topographic analysis of raw EC establishes fundamental signal propagation map for various visual conditions, characterizing the essential pathways along the visual hierarchy. We next sought to resolve how these signaling patterns are dynamically reconfigured for specialized visual processing by calculating the category-specific differential EC (e.g. ΔEC_others_). Using two pivotal hubs—the L-FFC and L-PHA2—as representative examples, we examined the distinct EC profiles underlying their task-selective neural activities to elucidate the signaling basis of their localized responses and reveal how these key regions orchestrate the integration of distributed regions into a coherent functional system. All connectivity modulations described hereafter reached strict statistical significance within the ΔEC matrix after Bonferroni correction (*p* < 3.5e^−7^; see Table S7 for L-FFC and Table S8 for L-PHA2).

During Face processing (vs. Others, Figure 6), the L-FFC functioned as a high-integration convergent hub. It received a massive surge of bottom-up in-flow from the ventrolateral stream, notably from L-PIT (*t* = 11.53), L-LO2 (*t* = 6.12), and bilateral V4 (*t* > 6.28). This sensory influx was complemented by top-down feedback from frontoparietal regions (e.g. L-IFSp, *t* = 6.67; bilateral LIPd, *t* > 6.15), while the L-FFC simultaneously augmented its out-flow to the bilateral Amygdala (*t* > 5.68), likely supporting the transfer of affective information. Internally within visual cortex, the L-FFC maintained robust bi-directional interaction with the L-PIT (*t_in_* = 11.53, *t_out_* = 8.98) and its contralateral counterpart R-FFC (*t_in_* = 9.88, *t_out_*_=_ 10.00). This specialized integration was balanced by a systemic suppression of the scene-processing stream, evidenced by significant down-regulation toward PHA1-3 and VMV1-3 (*t* < −6.76). When the task switched to Place processing (vs. Others, Figure 6), the L-FFC underwent a strategic functional reconfiguration. While its core face-selective pathways were suppressed (e.g., L-PIT: *t_in_* = −7.38, *t_out_* _=_ −8.52), the L-FFC significantly increased its out-flow toward the active ventromedial scene network, specifically targeting PHA1-3 (*t* > 11.14) and VMV1-3 (*t* > 8.72).

**Figure 6.**
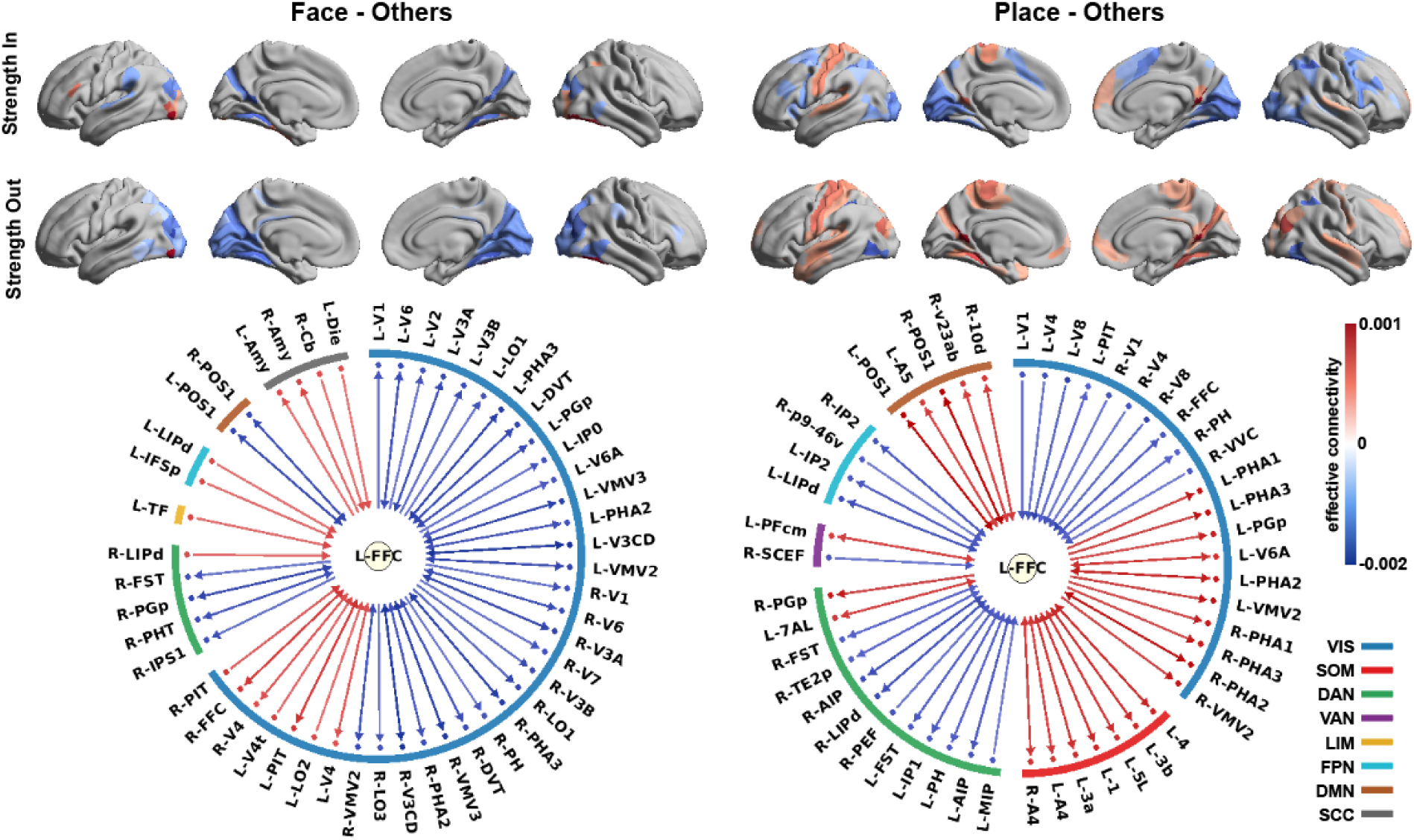
Reconfiguration of signaling patterns of category-selective hubs. This figure illustrates the task-specific effective connectivity profiles for the left fusiform face complex (L-FFC). The layout is organized into two primary sections: Face vs. Others (left) and Place vs. Others (right). Within each section, the top panels display the cortical topography of significantly ΔEC_others_ strength, separated into In-flow (first row) and Out-flow (second row). The bottom panels feature circular connectivity diagrams centered on the seed ROI. In these diagrams, arrows represent the direction of effective connectivity, with the color scale indicating the value of the ΔEC. The outermost ring is color-coded to denote the membership of target regions within the eight canonical functional networks. For visual clarity, only the top 50 significant edges are presented in the circular diagrams; the complete set of regulated connections can be found in Tables S7. All displayed edges reached statistical significance at *p* < 0.05 (Bonferroni corrected).

For L-PHA2, during Face processing (vs. Others, Figure S15), it was largely decoupled from the visual system, characterized by a systemic down-regulation of connectivity with both the core face network (L-FFC: *t_in_*= −11.37, *t_out_* = −6.07; L-PIT: *t_in_* = −10.18, *t_out_* = −7.38) and early-to-mid visual areas (L-V4: *t_in_*= −8.72, *t_out_* = −6.29; L-V8: *t_in_* = −8.11, *t_out_* = −5.99). This suppression highlights a competitive mechanism that minimizes cross-modal interference from the scene-processing machinery during face-related tasks. In contrast, during Place processing (vs. Others, Figure S15), the L-PHA2 acted as a primary orchestrator within the ventromedial stream. It received intensified inputs from the early visual cortex (L-V1/V2, *t* > 13.31) and middle-level visual areas (V4, *t* > 24.66; V8, *t* > 18.66), alongside higher-order modulation from L-IFSp (*t* = 11.15) and L-IPD/IP0 (*t* > 5.86). The L-PHA2 demonstrated extensive bi-directional coordination with adjacent regions, reinforcing a local spatial circuit with PHA1/3 (*t_in_* > 10.94, *t_out_* > 8.51) and VMV3 (*t_in_* > 13.34, *t_out_* > 12.70). These findings demonstrate that category-selective processing emerges from the dynamic reconfiguration of directed signaling, where the convergent functional hubs flexibly modulate their effective interactions with other regions to prioritize the influx of task-relevant information and suppressing competing streams to ensure coherent cognitive performance.

## Discussion

This study introduces a deep-learning-based framework for spatiotemporal modeling of task fMRI, providing, to our knowledge, the first comprehensive mapping of the hidden regulatory architecture that governs human visual processing at a whole-brain scale. Unlike GLM-based approaches that treat regional BOLD fluctuations in isolation, the STREAM framework not only provide the functional connectivity and activation map as traditional task-fMRI analysis, but also characterizes the whole-brain directed information flow by learning the latent transition functions of neural signaling. This flow-based perspective emphasizes how task demands perturb the brain’s signaling landscape, thereby driving the localized dynamic profiles observed across the visual hierarchy. Using this approach, we demonstrate that category-selective visual processing emerges from a fine-grained reconfiguration of directed signaling, where key hubs like the FFC flexibly switch their roles to meet specific cognitive demands. Our work thus establishes a scalable paradigm for uncovering the dynamical signaling principles that coordinate complex cognition across the human mind.

The success of STREAM highlights the transformative potential of artificial neural networks in capturing the non-linear, high-dimensional dynamics of the task-active brain^33^. Unlike traditional linear models or rigid causal frameworks like DCM^23^, deep learning networks effectively treats fMRI time series as continuous trajectories of a complex dynamical system. Previous research has demonstrated that such generative frameworks can successfully model resting-state dynamics and infer effective connectivity through virtual perturbations^28^. Our study successfully transferred this idea to modeling the dynamics of the task-active brain. By achieving exceptionally high spatial correlations with empirical GLM maps and faithfully reconstructing group-level functional topology, we demonstrate that deep learning models serve as a robust computational surrogate for neural population activity. It is important to note that while the MLP architecture emerged as the optimal model in our framework, it represents one of several viable solutions. Our comparative analysis reveals that Residual networks also exhibit remarkable stability and biological fidelity, suggesting that the capacity to learn underlying neural dynamics is a general strength of deep learning networks rather than a property restricted to a specific architecture. In contrast to studies that derive neural dynamics from predefined biophysical assumptions or structural connectivity priors^36,37^, STREAM prioritizes the consistency of dynamical evolution between network outputs and empirical fMRI signals, rather than enforcing mechanistic identity with the human brain^38^. Therefore, our primary objective was to establish the feasibility and validity of the deep learning-based perturbation inference framework, rather than constructing a brain-like network that mimics human neuroanatomy. Nevertheless, the fact that our models successfully recover hierarchical information gradients and task-specific pathways underscores the robustness of this data-driven approach. Future research incorporating structural constraints may further bridge this gap, yielding models that are both predictive and mechanistically interpretable^39^.

One of the most striking findings of our study is the strong spatial coupling between regional BOLD activations and incoming informational flow, which stands in stark contrast to the relative independence of outgoing influence. This distinction provides a potential computational explanation for recent evidence challenging the canonical interpretation of the BOLD signal^40^, which suggest that BOLD signal changes can frequently violate actual oxygen metabolism, a phenomenon particularly prevalent in the DMN. Our results imply that such discrepancies may stem from the inherent limitations of localized BOLD modeling where high BOLD activations faithfully reflect signal convergence and post-synaptic integration^30^, they are less indicative of the influence exerted by the regions originating these regulatory commands^29,30^. This divergence is topographically exemplified in visual category tasks, i.e., core In-flow regions, which align with GLM-based activations, are predominantly localized within the VOTC. In contrast, the primary Out-flow hubs reside within the DMN and FPN regions that may execute the top-down regulatory roles. This finding redefines the DMN, often dismissed as a passive or task-negative network, as a prominent source of outgoing influence that actively orchestrates task-related dynamics. Furthermore, we observed that even the intensity of down-regulated In-flow significantly correlates with GLM activation. This reinforces the physiological principle that the BOLD signal is a marker of the computational workload associated with total synaptic processing, regardless of whether the functional impact is excitatory or suppressive^29^.

By applying our framework to visual category processing, our results align with and extend the connectivity-constrained hypothesis^41,42^. We demonstrate that category-selective perception is not a fixed outcome of isolated modules, but emerges from the dynamic reconfiguration of effective connectivity^43^. While previous methodologies left the exact routes of such reconfiguration unclear, the STEAM framework allows us to resolve the fine-grained wiring patterns governing vision tasks. For instance, the FFC functions as a flexible computational resource that is strategically re-integrated into the ventromedial scene-processing stream when task demands shift. This macro-scale flexibility may be fundamentally underpinned by the rapid, concerted switching of neural codes within these regions, where neurons do not merely perform static specialized processing but instead undergo a temporal evolution of their functional roles^44^. Crucially, this selective integration is balanced by an active suppression mechanism, as evidenced by the systematic decoupling of PHA2 during face tasks to minimize cross-modal interference. Together, these findings establish that the visual hierarchy supports diverse cognitive functions through the flexible reconfiguration of pathways, providing a causal mechanism for how the brain balances specialized feature extraction with holistic integration. Despite these insights, several considerations warrant future exploration. First, while the STREAM framework effectively captures dynamics from fMRI, the inherent hemodynamic lag and low temporal resolution of the BOLD signal remain fundamental constraints. Although our model utilizes time-series modeling to infer directed interactions, the resulting effective connectivity primarily characterizes the spatial orchestration of information flow rather than the precise temporal progression of visual processing. Consequently, the millisecond-scale cascades of neural signaling that unfold immediately after stimulus onset remain largely obscured. Integrating this framework with high-temporal-resolution modalities, such as MEG or intracranial EEG, could provide a more granular view of how effective connectivity shifts unfold within the critical first few hundred milliseconds of perception. Second, while our group-level findings are robust, the observed inter-individual variability highlights the challenge of achieving stable causal inference at the single-subject level. Addressing this likely requires collecting extensive fMRI data for each participant to adequately capture their unique neural dynamical states. Finally, although our analysis covered the entire cortex, it was conducted at the parcellated ROI level, which inevitably sacrifices fine-grained spatial information. While fMRI offers the potential for voxel-wise mapping, scaling this deep-learning-based perturbation framework to tens of thousands of voxels entails significant computational costs. Developing more efficient architectures to enable high-resolution, voxel-level causal mapping represents a critical direction for future optimization. Consequently, while STREAM establishes fundamental principles of task-active signaling, future research is needed to refine it for more personalized and spatially exhaustive applications.

## Conclusion

In summary, we introduce a novel deep-learning-based framework for modeling task-fMRI time series, enabling the inference of causal signaling dynamics within the active brain. By applying this approach to visual category processing, we uncover a principle of dynamic reconfiguration that reconciles modular and distributed theories of perception. Looking forward, this framework provides a scalable and robust platform for investigating neural communication across diverse cognitive domains, serving as a critical methodological foundation for a more integrated and causal understanding of the human connectome.

## Data and code availability

The neuroimaging data used in this study were provided by the Human Connectome Project (HCP), and can be accessed through the ConnectomeDB platform (https://db.humanconnectome.org/).

A demonstration version of the custom code and a sample dataset are provided as Supplementary Information to facility peer review. The full, finalized version of the code and the STREAM framework will be made publicly available on GitHub upon formal publication.

## Acknowledgments

This work was supported by the National Natural Science Foundation of China (32427802, U24A200313, 32500926, 32400887), and the Key Research Development Project of Zhejiang Province (2026C02A1262(SD2)). Data were provided in part by the Human Connectome Project, the WU-Minn Consortium (principal investigators: David Van Essen and Kamil Ugurbil; 1U54MH091657) funded by the 16 NIH institutes and centers that support the NIH Blueprint for Neuroscience Research and by the McDonnell Center for Systems Neuroscience at Washington University.

## Methods

### Participants and Dataset

The MRI data were obtained from the Human Connectome Project (HCP) S1200 release^45^. From this dataset, we selected healthy young adults (age range: 22–35 years) who completed the full suite of task-fMRI sessions. All participants provided informed consent, and data collection was approved by the Institutional Review Board at Washington University in St. Louis. All data were acquired using a customized 3T Siemens Skyra scanner. Functional images were collected using a gradient-echo EPI sequence (TR = 720 ms; TE = 33.1 ms; flip angle = 52°; FOV = 208 × 180 mm; multiband factor = 8; voxel size = 2.0 mm isotropic).

We ensured high data quality without missing task blocks, resulting in a final sample of 1074 subjects for all subsequent analysis. Data were processed using the HCP Minimal Preprocessing Pipeline^46^, which includes gradient unwarping, motion correction, field map-based EPI distortion correction, and non-linear registration to the MNI space, followed by alignment to the standard CIFTI grayordinates space via the MSMSulc and MSMAll algorithms.

### Working Memory Task Paradigm

Participants performed a version of the N-back task, designed to map working memory and category-specific processing^47^. The task was presented in two runs, each containing blocks of four visual categories: faces, places, tools, and body parts. Each run included 8 task blocks (10 trials each, 25 s per block) and 4 fixation blocks (15 s each), covering both 0-back and 2-back conditions. For the purpose of neural dynamics modeling, we extracted the BOLD time series corresponding to all category-specific blocks, collapsing across memory loads to focus on category-selective signals. The fixation blocks served as the Baseline condition, providing a neural benchmark to evaluate the task-driven reconfiguration of the effective connectome.

### Parcellation and Time Series Extraction

To capture a whole-brain perspective, we parcellated the cortex into 360 regions (180 per hemisphere) using the HCP Multi-Modal Parcellation^48^, supplemented by 19 subcortical structures, totaling 379 regions of interest (ROIs). The representative BOLD time series for each ROI were extracted by averaging the signals across all constituent units. Following extraction, the signals were linearly detrended. For each subject, the category-specific time series were first preprocessed to remove low-frequency drifts using a high-pass filter (0.008 Hz) and linear detrending. the filtered signals were then standardized (z-score) to ensure consistent scaling. Following this global preprocessing, the time series were partitioned according to the experimental design matrix. Specifically, BOLD segments corresponding to each task condition (e.g., Face, Place) were extracted from their respective blocks and subsequently concatenated to form condition-specific time series. This procedure ensured that the modeling of individual-level neural dynamics was focused on the steady-state signaling patterns unique to each cognitive demand, while maintaining the temporal continuity necessary for learning latent transition functions.

### Model Selection and Refinement

To identify the most capable architecture for representing task-driven neural dynamics, we first conducted a benchmarking study on a subset of 100 subjects from the HCP. We compared five candidate architectures^28,38^: Vector Autoregression (VAR), Multilayer Perceptron (MLP), Recurrent Neural Networks (RNN), Long Short-Term Memory (LSTM), and Residual Networks (ResNet). Each model was initialized based on established parameters to predict the state of 379 cortical regions at time t+1 using a three-time-point historical window (t-2, t-1, and t). Models were evaluated based on predictive robustness (test-set MSE) and biological fidelity.

The latter was assessed by evaluating the model’s capacity to autonomously generate realistic neural dynamics. To initialize the generative process, the input window was seeded with three initial time points. To break potential symmetry and simulate physiological variability, the seed was augmented with a small amount of Gaussian noise (sigma = 0.1). Following this initialization, the model then iteratively predicted the subsequent neural state at t+1, which was recursively fed back as the input for the next time step. Through this recursive process, we generated a simulated time series of 160 time points, a duration chosen to match the length of the empirical task blocks used during training (∼153 TRs). To ensure biological fidelity, we calculated the model-derived functional connectivity (FC) using Pearson correlation coefficients between the simulated time courses of all ROI pairs. This was then compared against the empirical task-FC, computed via the same Pearson correlation analysis on the concatenated, condition-specific empirical signals.

The MLP architecture was selected as the core model due to its superior performance during benchmarking. We observed that while parameters optimized for resting-state dynamics provided a baseline, they required systematic refinement for task-modulated BOLD signals. We first optimized the prediction horizon by evaluating length of the historical time window ranging from one to five TRs, identifying a length of 3 TRs as the optimal balance between instantaneous accuracy and stable FC reconstruction. To further enhance the model’s representational capacity, we implemented a deep architecture consisting of three hidden layers with a Hidden Dimension Multiplier of 2.0 and a Latent Dimension Multiplier of 1.0 relative to the number of ROIs. The framework was trained using the Adam optimizer with a learning rate of 2 × 10^−3^ and L2 regularization (weight decay = 5 × 10^−5^). Training was conducted over 120 epochs with a batch size of 50, incorporating a ReduceLROnPlateau scheduler to ensure convergence. To prevent overfitting and promote generalization across the extensive HCP cohort, we integrated Dropout (rate = 0.1) and standardized the inputs to unit variance. This specialized deep-learning configuration, which we termed STREAM (Spatiotemporal Representation for Effective connectivity Analysis Model), established a robust computational framework for mapping neural communication, providing a scalable platform for investigating the spatiotemporal principles of task-modulated brain states.

### Comprehensive Validation of the STREAM framework

To systematically ensure that STREAM faithfully captured task-driven neural dynamics, we performed a multi-dimensional evaluation of model performance. Building upon the iterative generation process described above, we extended the synthetic BOLD signals to a duration of 160 time points for each task condition. We first assessed the model’s capacity to reconstruct macro-scale functional architectures by spatially correlating the group-averaged “model-derived FC” with the empirical group-level task-FC. Secondly, to ensure the generalizability of the learned transition rules, we conducted a split-half reliability analysis. The full cohort (N = 1074) was randomly divided into two independent subgroups; the spatial correlation between their respective model-derived FC matrices was quantified to evaluate the stability of the framework across different subject pools. Finally, we verified the biological grounding of STREAM by examining whether its generative dynamics reflected empirical brain activations. For each of the 379 regions, task-specific activation was quantified as the mean of the simulated time series during task conditions relative to the simulated baseline state. These model-derived activation profiles were then compared with empirical task-contrast maps (quantified as Cohen’s d effect sizes). This cross-modal validation confirms that the latent representations learned by STREAM are not merely statistical fits but are deeply aligned with established neurobiological patterns.

### Inference of Effective Connectivity

To quantify the directed interaction between brain regions, we employed the Neural Perturbational Inference (NPI) approach^28^. Rather than relying on traditional cross-correlations, NPI infers effective connectivity by measuring the causal response of the trained network to systematic perturbations. To derive a robust estimation of task-specific EC, we performed the perturbational analysis across all available temporal contexts within the empirical task sequence. For each subject and condition, we extracted every valid historical window of 3 TRs (the “seed” states) directly from the empirical time series. For each window, we applied a unit perturbation to a single source region i at the final time point (t) and calculated the resulting change in the predicted activity of all target regions at time t+1, relative to the unperturbed model output. The final EC weight from region i to region j was defined as the average causal response across all empirical time windows for that specific task condition. This perturbational approach inherently produces an asymmetric EC matrix, as the causal influence of region i on region j is not mathematically or biologically identical to the influence of j on i. By repeating this process for all 379 regions, we constructed a full directed EC adjacency matrix for each subject. Finally, these individual matrices were averaged across the cohort to derive a representative group-level EC map. To characterize the directional topography, the EC matrix was further decomposed into in-flow (the sum of incoming causal influences to a region) and out-flow (the sum of outgoing causal influences from a region) metrics.

### Task- Driven Regulation of Effective Connectivity and Joint Analysis with Local Activation

To isolate the specific neural pathways regulated by distinct visual categories, we employed a subtractive logic to derive differential EC (ΔEC) matrices. For each participant, ΔEC was calculated by subtracting the baseline EC (defined by inter-block fixation periods) from the task-specific EC (e.g., Face, Place, Body, or Tool). This subtractive approach effectively extracted task-induced regulation in directed signaling from the brain’s intrinsic architecture. The ΔEC was further decomposed into up-regulated (positive ΔEC) and down-regulated (negative ΔEC) components. We quantified regional modulation intensity by calculating the sum of increased/decreased In-flow and Out-flow for each of the 379 regions.

To investigate the biological basis of these causal reconfigurations, we performed a joint coupling analysis with empirical task-activation maps (Cohen’s d). Given the distinct metabolic and signaling profiles of activated versus deactivated ensembles, we implemented a stratified spatial correlation analysis. Brain regions were partitioned based on the polarity of their GLM-based activation. Within each partition, we systematically correlated the Cohen’s d values with our four EC-derived regulation metrics (Positive/Negative In-flow/Out-flow). To ensure the reliability of these regulatory signatures, we performed paired t-tests (Task vs. Baseline) across the full cohort, applying a strict Bonferroni correction, with the significance threshold set at 0.05 / (379 × 378) ∼ 3.5e^−7^. This allowed us to repeat the coupling analysis using only statistically significant edges, thereby confirming the robustness of the in-flow-activation coupling.

### Systems-Level Statistical Analysis

To uncover the systems-level logic governing task-induced modulations, we aggregated regulatory intensities across eight functional sub-networks. Brain regions were partitioned by mapping the 379 ROIs of the HCP-MMP parcellation onto the seven canonical functional networks ^35^, supplemented by one subcortical module. This mapping provided a standardized framework to transition from region-specific connectivity to broader systemic interactions, encompassing the Visual (VIS), Somatomotor (SOM), Dorsal Attention (DAN), Ventral Attention (VAN), Limbic (LIM), Frontoparietal (FPN), Default Mode (DMN), and Subcortical (SCC) networks.

To ensure the statistical validity of the network-level comparisons, we employed a group-to-individual back-projection approach. For each task condition, we first established a “group-level regulatory template” by identifying the set of edges that exhibited significant modulation (e.g., all edges showing significantly up-regulated In-flow at the group level). Using these templates as spatial masks, we then extracted the corresponding average regulation intensity from each individual’s EC matrix for every sub-network. This procedure ensured that the statistical comparisons were performed on consistent edges across all participants.

### Topographic Mapping of Recurrent Hierarchical Information Flow

To characterize the dynamic routes of information propagation across the visual hierarchy, we prioritized the analysis of the ventral visual stream, which serves as the primary pathway for visual category processing. We constructed a comprehensive ventral sub-network by extracting bilateral regions of interest (ROIs) from the HCP Multi-Modal Parcellation, ensuring the selection meticulously encompassed the full functional trajectory of the ventral pathway. This anatomical definition included early visual areas from V1 through V4 (incorporating subsets V3A/B, V3CD, and V4t), mid-level motion and object-related regions such as MT, MST, FST, V8, and LO1–3, and the high-level ventral temporal areas specialized for category recognition, including the posterior inferotemporal cortex (PIT), fusiform face complex (FFC), ventral medial visual areas (VMV1–3), parahippocampal areas (PHA1–3), and the ventral visual complex (VVC).

To isolate the most robust and biologically plausible pathways of information transfer within this extensive stream, we implemented a dual-directional “Top-K (K = 5)” backbone extraction algorithm. For each node within the ventral sub-network, we retained only the 5 strongest incoming and 5 strongest outgoing connections based on their effective connectivity (EC) magnitudes. This pruning strategy effectively filtered out weak or non-essential interactions to reveal the core causal skeleton of task-modulated signaling. Within this pruned architecture, we quantified the hierarchical position of each region by defining an effective path cost for each directed edge, calculated as the inverse of the effective connectivity magnitude (1/|EC|). Using Dijkstra’s shortest-path algorithm^49^, we computed the cumulative the “informational distance” from the primary source in the left V1 to all subsequent regions in the stream. This structure was visualized using a force-directed layout (Figure 5). In this representation, the x-axis represents “distance” from the source, quantifying the cumulative cost of information transfer. The y-axis represents the “Out-flow Intensity,” defined as the sum of a region’s outgoing effective connectivity, which characterizes its capacity to broadcast signals to downstream targets.

### Seed-Based Analysis of Task-Specific Connectional Reconfiguration

To investigate how category-specific processing emerges from the dynamic reconfiguration of neural interaction patterns, we performed a seed-based analysis on the effective connectivity of two pivotal hubs, namely the L-FFC for face processing and the L-PHA2 for place processing. We utilized the ΔEC matrices derived by contrasting each specific task condition against the average of all other categories to subtract the common sensory-motor and intrinsic signaling background. This contrast allowed us to isolate the reconfiguration uniquely dedicated to specialized category processing.

For each seed region, we characterized its task-driven reconfiguration by identifying and quantifying all path-specific regulations that reached statistical significance under both matching and non-matching task conditions. Specifically, we examined the L-FFC during face processing as the matching condition and during place processing as the non-matching condition, with a reciprocal analysis performed for the L-PHA2. For each condition, we conducted paired t-tests across the cohort for all incoming and outgoing connections associated with the seed, applying a strict Bonferroni correction [p < 0.05 / (379*378)] to ensure the robustness of the detected modulations.

